# MPRAudit Quantifies the Fraction of Variance Described by Unknown Features in Massively Parallel Reporter Assays

**DOI:** 10.1101/2020.02.12.945113

**Authors:** David A. Siegel, Olivier Le Tonqueze, Anne Biton, David J. Erle, Noah Zaitlen

## Abstract

Transformative advances in molecular technologies, such as massively parallel reporter assays (MPRAs) and CRISPR screens, can efficiently characterize the effects of genetic and genomic variation on cellular phenotypes. Analysis approaches to date have focused on identifying individual genomic regions or genetic variants that perturb a phenotype of interest. In this work, we develop a wholistic framework (MPRAudit) to determine the global contribution of sequence to phenotypic variation across subsets of the entire experiment, opening the door to myriad novel analyses. For example, MPRAudit can reliably estimate the upper limit of predictive performance, the fraction of variation attributed to specific biological categories, and the total contribution of experimental noise. We demonstrate through simulation and application to several types of real MPRA data sets how MPRAudit can lead to an improved understanding of experimental quality, molecular biology, and guide future research. Applying MPRAudit to real MPRA data, we observe that sequence variation is the primary driver of outcome variability, but that known biological categories explain only a fraction of this variance. We conclude that our understanding of how sequence variation impacts phenotype, even at the level of MPRAs, remains open to further scientific discovery.

## Introduction

Massively parallel reporter assays (MPRAs) have transformed our ability to directly investigate the effects of genetic perturbations on cellular phenotypes such as gene expression, growth, and chemical resistance. They allow researchers to test tens of thousands of sequences at once, which provides vast amounts of data to study complex phenomena while minimizing cost and the impact of batch effects. MPRAs are now a fundamental tool and have been used to identify enhancer activity[1], determine the influence of chromatin on cis-regulatory sequences [2], identify sources of variation in red blood cell traits [3], and identify functional annotations in 3′ untranslated regions[4]. Related assays such as CRISPR screens have identified loss-of-function mutations affecting tumor growth and metastasis in mice[5] and genes that regulate T cell activation in the genome [6, 7]. This type of screening can also be used for drug discovery and the identification cytotoxic compounds [8, 9].

Several computational and statistical methods have been designed specifically for the analysis of MPRAs, focusing on methods for differential expression[10, 11, 12, 13, 14, 15, 16]. Myint et al [10] classified MPRAs into three broad categories: characterization studies[17, 18, 19], which examine and classify the sequence features of regulatory elements; saturation mutagenesis studies[4, 20, 21, 22], which look at the impact of all possible mutations to a functional element; and differential analysis studies[23, 24], which seek to determine the differential impact of multiple variants. In a fourth category, prediction studies, no differential analysis is performed, but a large number of sequences or features are measured and an out of sample predictor is created [25, 26, 27]. We collectively term these approaches as bottom-up: the analysis objective is to identify which individual perturbations have a causal impact on the phenotype of interest.

In this work we develop a top-down MPRA analysis framework: the analysis objective is to determine the contribution of all sequence variation to experimental outcomes across the entire experiment. MPRAudit takes as input annotated subsets of the MPRA experiment and outputs estimates of the biological and technical variance across these subsets. These parameters have a range of utilities across MRPA use cases, and can be applied even when there is not sufficient power to identify individual causal perturbations with bottom-up approaches. For studies that consider perturbations to individual elements, MPRAudit provides an estimate of the quality of the dataset and sets an upper limit to the performance of prediction assays. For studies that consider pairs of elements with and without a mutation, MPRAudit can additionally determine the contribution of the flanking sequence of a mutation to the effect size of that mutation. For studies that consider groups of elements, MPRAudit tells researchers how much novel biology is left once known biological classes are accounted for; and what subsets contain unknown biological sources of variation waiting to be discovered.

We evaluate MPRAudit through application to simulated and real data sets. In realistic simulations, we show that MPRAudit gives accurate estimates of the variance explained by technical factors, and that MPRAudit’s uncertainty estimates accurately match the empirical variation across simulations. We then apply MPRAudit to a simulated MPRA experiment, where we demonstrate the use of MPRAudit on simulated data of groups of gene expression measurements, paired effect size measurements, or groups of measurements. Finally, we apply MPRAudit to experimental data, including our recent MPRA on the 3′ UTR in human cell lines, where we demonstrate the use of MPRAudit on categorical data and show that the cis-regulatory effects of the 3′ UTR sequences on gene expression remain largely unexplained by available models.

MPRAudit is implemented as a free and open source software package. MPRAudit can be used in any barcoded molecular assay, as it is based on holding out or resampling over barcoded clones within a dataset and it does not require parametric model assumptions.

## Results

### Overview of Idea

#### Definitions

We define an MPRA as an experiment with an outcome *O*_*i*_ for each of *N* perturbations, *i* ∈ {1…*N*}, where each outcome has multiple clonal replicates (“clones”) *c* ∈ {1…*n*_*j*_} (*n*_*j*_ might vary with *i*). In many cases, the perturbations are sequences and the outcomes are the aggregated count ratios of RNA 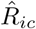 to DNA 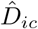. In this setting:

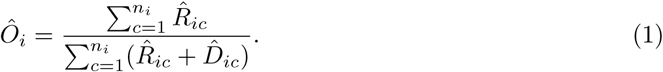

In other studies *O*_*i*_ might be a log-ratio, or a ratio of ratios, and these can be directly incorporated into MPRAudit. The perturbations do not need to be sequences with RNA/DNA outcomes; they could be deletions in a CRISPR screen, or chemicals in a drug discovery assay.

We index unique DNA sequences with *S*_*i*_. The clonal replicates must differ by only a barcode, which is a common practice in MPRAs [28] because it effectively provides replicates allowing researchers to compute p-values and confidence intervals. MRPAudit uses the clonal replicates to obtain estimates of the uncertainty in aggregated outcomes.

We assume each measurement *Ô*_*i*_ is generated from:

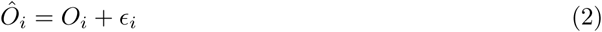

where *ϵ*_*i*_ is noise. The major advancement in our manuscript is the ability to calculate *var*(*ϵ*_*i*_|*i*), which can be done in great generality by performing a statistical jackknife over the clones that make up the statistic in Eq. 1 (see Methods). This provides direct estimates of the contribution of noise to an experiment. The quantity that we focus on is the fraction of variance in our measurements that is determined by sequence variation, which we call the “explainability” and denote by *b*^2^ (by analogy with the heritability *h*^2^ in genetic studies or *r*^2^ in regression studies):

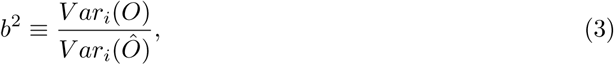

where the subscript i denotes summation over sequences: 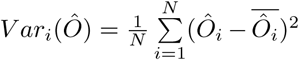. The under-lying principle is that the variability in repeated measurements of identical clones can provide us with an estimate of *V ar*(*ϵ*_*i*_|*i*) for each sequence [29]. If the measured variability across sequence outcomes is significantly greater than what can be ascribed to the barcoded sources of variation, then in the absence of clonal confounders the remainder must be due to sequence variation.

MPRAudit captures all of the variability from clone to clone and compares it to the variability in outcome across perturbations. On one extreme, there might be an experiment where the clones from the same perturbation produce highly variable results. In that case *b*^2^ would be low and the “technical” sources of variation would be large in comparison with the “sequence-based” variability. On the other extreme, there might be an experiment where clones produce nearly identical outcomes, but the perturbations each differ substantially from each other. In that case *b*^2^ would be high and the technical sources of variation would be small in comparison with the sequence-based variability.

*b*^2^ depends on both technical parameters like sequencing depth, as well as the experimental design and has many desirable features for top-down MPRA analyses: If the sequencing depth is increased, then *b*^2^ will increase. If the sites chosen for perturbation are not functional, then *b*^2^ will be 0. The nonparametric nature of MPRAudit means that *b*^2^ can be accurately estimated even under non-normal noise distributions for *ϵ*_*i*_.

#### Technical and Biological Factors in an MPRA

There are many technical and biological factors that can drive the variability from clone to clone and from sequence to sequence in an MPRA.

At the clone level, there are several technical factors that might drive the variability from clone to clone. For MPRAs, one is the sequencing read depth, which has been observed to follow a negative binomial distribution across clones[15]. Another is PCR, which is known to be nonlinear in some regimes[30, 31].

There are also sources of biological variability that drive differences in RNA concentration from clone to clone. The number of RNA molecules in a cell varies due to cell-cycle and other biological sources of variation. Clonal cells lines have some genetic and epigenetic variability between seemingly identical clones[32]. In fast-UTR[4] and related assays, the site at which the DNA is inserted into the host genome can also play an important role in gene expression.

At the sequence level, differences in sequence (such as GC-content) might also lead to systematic bias in the amplification and sequencing steps, which would have to be accounted for separately.

## Simulation Setup and Results

### Read Count Simulation Setup

In order to verify the accuracy of MPRAudit, we first examine a simulation based on our previously developed MPRA called fast-UTR[4]. In a fast-UTR experiment, a large set of test sequences are inserted into the 3’ UTR of a reporter transgene in order to determine the effects of these sequences on gene expression (RNA stability and/or steady state RNA levels). Each sequence is replicated multiple times in the reporter library, and replicate clones can be distinguished by the incorporation of distinct nucleic acid sequences (clone indices). After introduction of the reporter library into a population of cells, amplified RNA and DNA molecules are sequenced with the expectation that the read counts will be roughly proportional to the average numbers of RNA and DNA molecules per cell. Effects of sequences on steady state RNA levels are determined by comparing RNA read counts to DNA read counts in cells at steady state (t0). Effects on RNA stability are determined by treating cells with doxycycline, which prevents transgene transcription, and comparing RNA levels before and after (e.g., 4 h after, *t*_4_) addition of doxycycline.

In our simulation, we use *i* ∈ {1…*N*} as sequence indices and *c* ∈ {1…*n*_*i*_} as clone indices. We begin our simulation by selecting some starting number of cells, and therefore some true number of DNA molecules, for each of *n*_*i*_ clones:

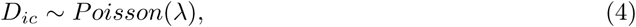

where *λ* is a constant that denotes the average number of DNA molecules per clone, and if the Poisson distribution yields zero counts of DNA we set the counts of DNA to 1. The variability in the number of cells per clone is what prompted us to obtain DNA counts in addition to RNA counts in our real MPRA experiment, since RNA counts are dependent on the number of cells per clone but the RNA/DNA ratio is not.

We then draw the true number of RNA molecules for each of those clones: We assume some RNA/DNA ratio ‘*A*_*i*_’ that is fixed for each clone in sequence *i*, but might vary from sequence to sequence; and we assume some normally distributed (biological) noise with variance proportional to *D*_*ic*_ and a noise parameter *s*^2^:

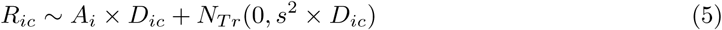

where 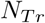 is a truncated normal (the number of RNA molecules must be nonnegative), and rounding is performed to obtain an integer number of RNA molecules. *s* represents a biological source of noise, such as cell-cycle, that will remain in the data in the limit of infinite sequencing, but can be accounted for as the number of clones becomes infinite.

*A*_*i*_ introduces sequence-based variation into the simulation. If *A*_*i*_ is constant across all sequences and *s* = 0, then the sequence-based variation is zero, so we expect the total variation to equal the technical variation and to find *b*^2^ = 0. If *A*_*i*_ varies significantly with sequence (i.e. *i*), then the sequence-based variation will be nonzero and the total variation will differ from the technical variation. In real experiments there might be additional forms of sequence-based variation from the PCR process (due to GC content, for instance), which we do not include in this simulation, but we do account for in the real analysis.

Given these numbers of simulated RNA and DNA molecules, simulated counts of RNA and DNA for each clone and sequence are drawn from a negative binomial distribution to simulate overdispersion. Further details are given in the methods section. The heavy tails, truncated distributions, and complex relationships between clonal counts and aggregated outcomes in these types of datasets would pose problems for parametric methods, which is why we favor the flexibility and accuracy of a resampling-based approach.

### Setup of Simulated Experiment

To test MPRAudit we have created a simulated experiment based on fast-UTR by simulating RNA and DNA read counts as described above, holding all parameters constant except varying *A*_*i*_ (the RNA/DNA ratio) from sequence to sequence.

We first create a simulated experiment by generating 10,000 sequences *S*_*i*_ of length 15 nucleotides (nt). The first 5,000 sequences are randomly generated with a uniform probability of A, C, T, or G at every location along the 15 nt. The second 5,000 sequences are identical to the first 5,000, except the sequence “TATACAG” has been substituted at a random location so that we can treat it as a fictitious miRNA binding site. For example, *S*_2_ might be GAGACAATGGATCAA, and *S*_5002_ might be GAG<u>TATACAG</u>ATCAA. Analyzing the difference in gene expression between *S*_2_ and *S*_5002_ provides information about the importance of the “TATACAG” motif to gene expression.

We then require these sequences to determine the RNA/DNA ratio (*A*_*i*_ in equation (5)) according to an additive linear model:

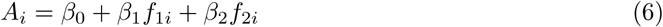

where *f*_*ki*_ is the number of times the *k*^*th*^ feature appears in the sequence *S*_*i*_, and *β*_*k*_ gives the effect size of *f*_*ki*_ on gene expression (*A*_*i*_). We set *f*_1*i*_ to be the number of times “TATACAG” appears in sequence *S*_*i*_, and *f*_2*i*_ to be the number of times “T” appears in the sequence *S*_*i*_. This choice of feature 2 implies that the nature of the TATACAG mutation will have a nontrivial affect on gene expression (for instance mutating “TATACAG” to “AAAAAAA” will have a different effect size than mutating “TATACAG” to “AAAAAAT”).

We set *β*_0_ = 2, and in a null simulation (where features have no effect on gene expression) we assume *β*_1_ = *β*_2_ = 0. To test the effect of sequence variation, we set *β*_1_ = −0.5 and *β*_2_ = 0.15. In this case each occurrence of “TATACAG” decreases gene expression and each “T” increases gene expression. Under the null they have no effect.

We performed the simulated experiment 1000 times and compare our estimates of the mean and standard deviation to their empirical values. We chose model parameters (including sufficient sequencing and clonal noise) such that the sequence-based variation makes up a significant fraction of the total variance. See methods for further details.

### Analysis Setup

The output of fast-UTR experiments is often given by the ratio:

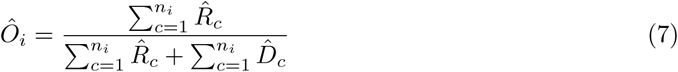

We have also implemented and tested MPRAudit with log(RNA/DNA), and we report some results with this choice of statistic in later sections of the manuscript.

For each simulation we use MPRAudit to calculate the explainability *b*^2^. To verify the accuracy of MPRAudit, we run the simulation with and without noise to perform a direct empirical measurement of Var(*O*) and Var(*Ô*), respectively, of equation (3). We then compare the results of MPRAudit to the direct calculation of *b*^2^ from equation (3). To run the simulation without noise, we set s=0 and the number of counts *K*_*R*_ and *K*_*D*_ to be very large (increasing both by a factor of 10^6^), keeping all other parameters fixed.

The empirical variance or standard deviation is obtained by running the simulated experiment 1000 times, calculating and recording the *b*^2^ each time, then calculating the standard deviation of the recorded *b*^2^ across the 1000 experiments at the end.

### Simulation Results for Single Perturbations

One of the primary use cases that we envision for MPRAudit is to quantify the total fraction of sequence variation across the entire experiment. The explainability *b*^2^ sets an upper limit on the accuracy of prediction methods, and provides a measurement of the reproducibility of the dataset and experiment as a whole. In addition, when a dataset is divided into groups (for instance by the presence or absence of a binding site motif), a comparison of *b*^2^ across groups determines the remaining biological signal left in each group, which could be informative for follow-up investigations.

We first simulate the null by setting *A*_*i*_ = 2 to be constant for all sequences, guaranteeing that *b*^2^ = 0 and that sequence will have no effect. The results shown in Figure 2 demonstrate that the bias is uniformly small, and rapidly decreases as the number of clones per sequence increases. We recommend using at least four clones per sequence to minimize this source of bias. Negative estimates of *b*^2^ are possible if the estimate of the technical variance is large, which could happen by chance if b^2^ is small – especially because this inflates jackknife estimates[33], which causes b^2^ to be biased downwards. To minimize this source of bias, when we jackknife over clones we use a delete-D jackknife rather than a leave-out-out jackknife[34]. The standard deviation of *b*^2^ predicted by MPRAudit matches the empirically determined standard deviation across simulations. Consistent with expectations, the standard deviation decreases as the number of clones increases.

**Figure 1:**
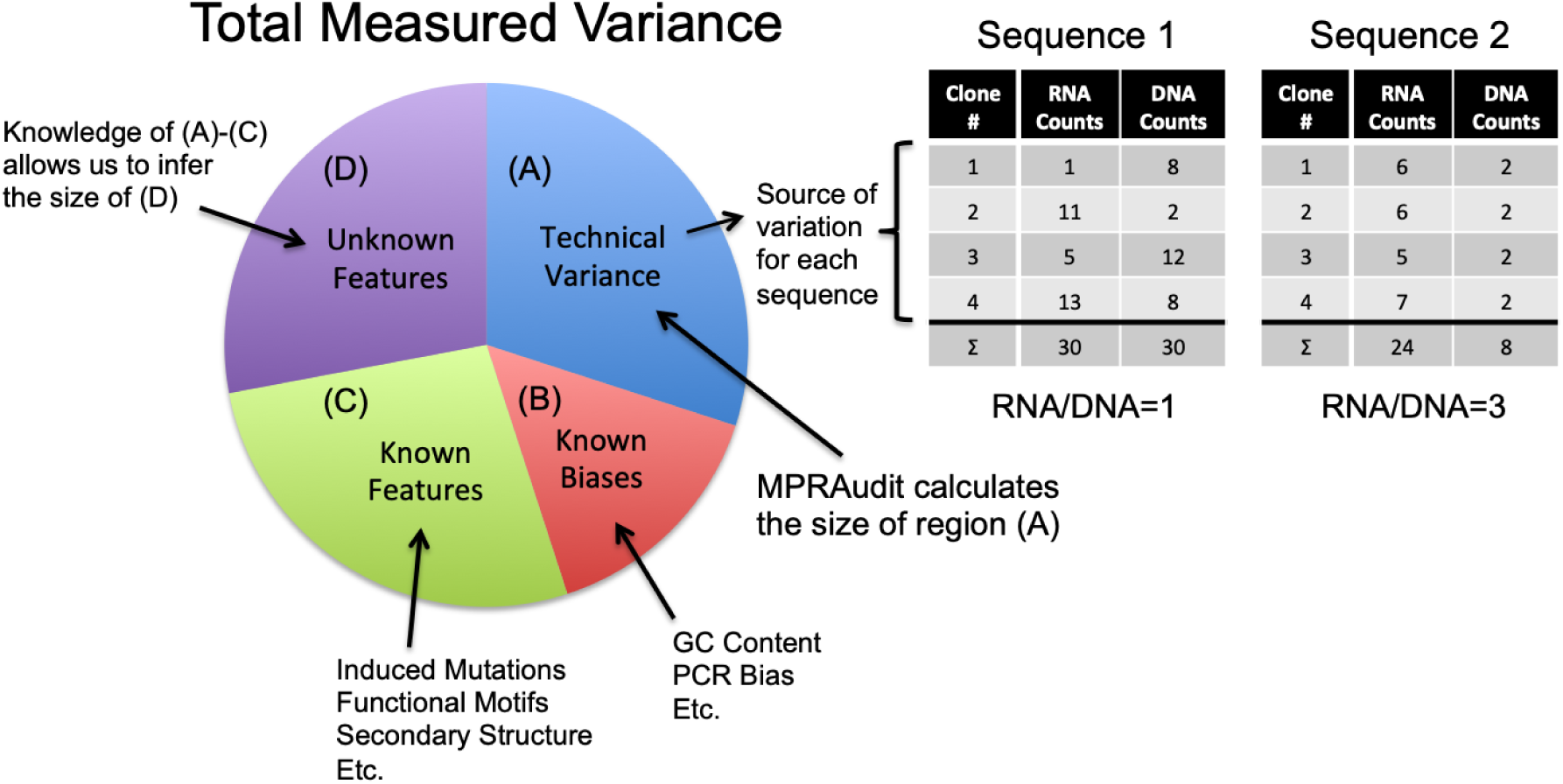
The total measured variance of any MPRA can be partitioned into technical (A) and sequence-based (B-D) sources of variation. MPRAudit estimates the technical variance from the variation from clone to clone for each sequence. In the examples given, the measured RNA/DNA ratio of Sequence 1 differs from Sequence 2, but the variance of Sequence 1 is much larger than the variance of Sequence 2. Sequence-based variation can be partitioned into sub-groupings: (B) sources of systematic bias, such as GC content; (C) known features; or (D) unknown features. The amount of sequence variation described by unknown features can be inferred from knowledge of (A)-(C). *b*^2^ is the fraction of variance explained by groups (B)-(D) out of the total variance.

**Figure 2:**
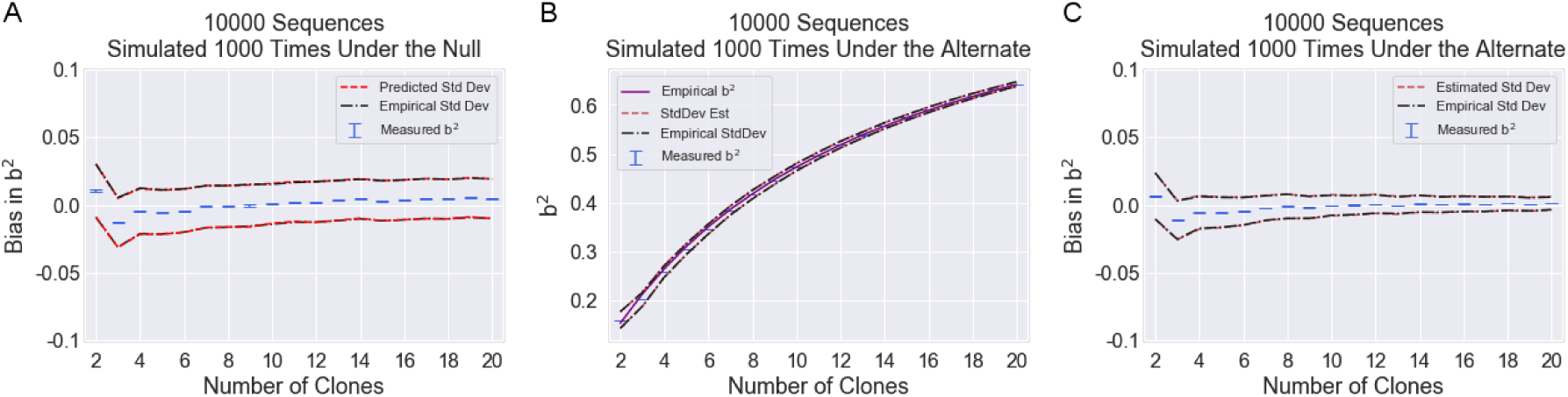
Results of explainability (b^2^) calculations for single perturbations. Error bars in blue (very narrow on this scale) give the standard error of b^2^ for the full 10,000 sequences. Dashed lines give estimated and empirical confidence intervals (standard deviation). (a) *b*^2^ for single perturbation data simulated under the null (*A*_*i*_ = 2, a constant), where *b*^2^ is zero. (b) *b*^2^ for data simulated under the alternate (*A*_*i*_ = 2 −0.5*f*_1*i*_ + 0.15*f*_2*i*_). The empirical value of b^2^ was calculated by running the simulation with and without noise. (c) Same data as (b), except the y-axis is given as bias in b^2^ (estimate minus empirical) instead of explainability.

To incorporate a biological source of variability, we set *A*_*i*_ = 2 − 0.5*f*_1*i*_ + 0.15*f*_2*i*_ and use MPRAudit to compute *b*^2^ for each simulation. We also compute the true *b*^2^ by running the same simulation with and without noise. Figure 2b shows that the *b*^2^ estimated by MPRAudit is a close match to this empirical value. *b*^2^ increases as a function of the number of clones in Figure 2b due to the decreased statistical noise. The bias is shown in Figure 2c to be uniformly small and rapidly decreases as the number of clones per sequence increases. In summary, Figure 2 shows that MPRAudit can accurately calculate the overall fraction of phenotypic variation due to sequence.

### Simulation Results for Groups of Perturbations

MPRAudit can also be used to quantify how much sequence variability remains after accounting for a known biological phenomenon. For example, if the binding site TATACAG is known, MPRAudit can use the same technology to calculate the within-group explainability of the grouped sequences to estimate the remaining biological explainability after TATACAG is accounted for. Details are given in the methods section.

We first simulate the null model where *A*_*i*_ is constant, and we expect the within-group *b*^2^ to equal zero since there is no biological variation. Figure 3a shows that MPRAudit gives an accurate estimate of *b*^2^ in this case with little bias, which decreases as the number of clones per sequence increases. It also gives an accurate estimate of the standard error, which decreases with the number of clones per sequence as well.

**Figure 3:**
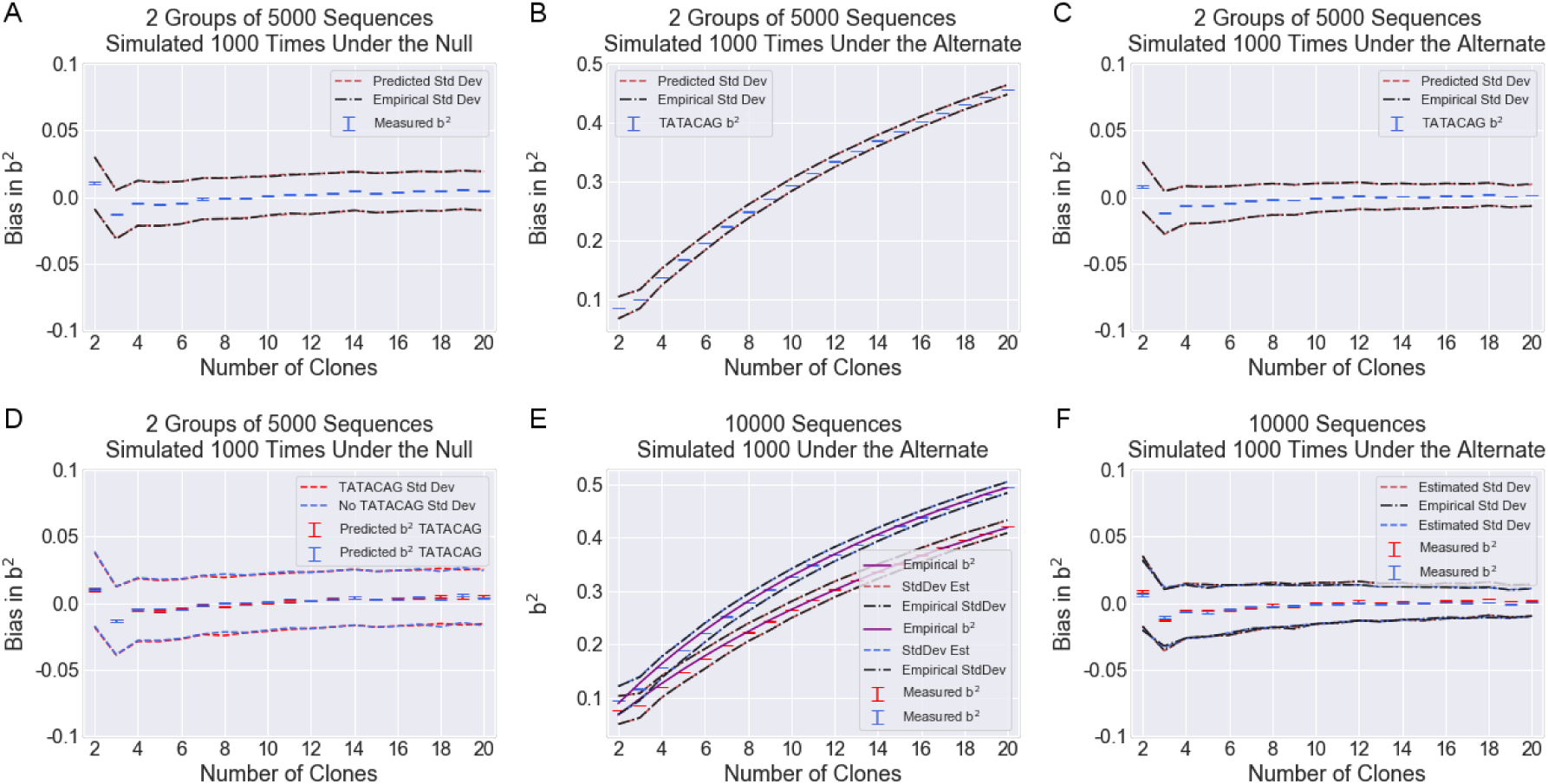
Results of explainability (b^2^) calculations for groups of sequences. We generate two groups of sequences in our simulation, those with TATACAG motifs and those without, with variation within groups corresponding to the number of T’s. Error bars (very narrow on this scale) give the standard error of *b*^2^ for the full 10,000 sequences. Dashed lines give estimated and empirical confidence intervals (standard deviation). (a) Total remaining *b*^2^ after TATACAG categories are accounted for under the null, where *b*^2^ is zero. (b) Remaining *b*^2^ after TATACAG categories are accounted for under the alternate, where sequences variation persists in correspondence with the number of Ts. (c) Same as (b), except the y-axis is given as bias in *b*^2^ (estimate minus empirical). (d) Explainability within each category of sequence (with or without TATACAG) under the null, where b^2^ is zero, (e) under the alternate, where there is less variability in the TATACAG group, (f) bias under the alternate.

We next study the effect of biological variability. We again set *A*_*i*_ = 2 − 0.5*f*_1*i*_ + 0.15*f*_2*i*_ and group sequences by the presence or absence of TATACAG. In this case we expect there to be some biological variation remaining within the groups, since the number of T’s in the sequence varies from sequence to sequence in both groups and has an effect on gene expression. Figure 3b shows that the *b*^2^ estimated by MPRAudit is a close match to the empirical value, while Figure 3c shows that the bias is small and rapidly decreases as the number of clones per sequence increases. MPRAudit’s estimated standard deviation once again agrees with the empirical value.

The set-based tests implemented by MPRAudit can determine whether unknown biological features are responsible for phenotypic variation, and can further identify which groups of perturbations are enriched for these unknown categories by calculating the *b*^2^ within each group. For the dataset simulated here, we expect the *b*^2^ within the 5,000 “TATACAG” sequences to be lower than the *b*^2^ within the 5,000 completely random sequences, since the TATACAG region has no variation. This is what we observe in Figure 3. In Figure 3e, the data is simulated under the null and there is no *b*^2^ in either group. In Figure 3f, the data is simulated under the alternate with *A*_*i*_ = 2 − 0.5*f*_1*i*_ + 0.15*f*_2*i*_, and *b*^2^ is shown to be higher for the group with completely random nucleotides. In Figure 3g the bias within each subgroup is shown to be similarly small.

### Simulation Results for Pairs of Perturbations (Effect Sizes of Mutations)

Another common use of MPRAs is to explore the effect of directed mutations inside specific regions of the genome. We define the effect size of a mutation to be the signed difference in gene expression between a sequence and its designed mutation (we use *P*_*ij*_ = *O*_*i*_ − *O*_*j*_ but one could alternatively measure fold change), and calculate the *b*^2^ of this quantity by applying the same technology that was developed for single perturbations to a slightly different setting. The details are discussed in the Methods section. To demonstrate this in a simulated experiment, we consider the 5,000 sequences with TATACAG to be wild-type (WT) sequences, and the 5,000 sequences with random mutations of the “TATACAG” motif to be their paired mutations (MT). This establishes 5,000 WT-MT pairs.

To study the null, we examine the effect size of TATACAG mutations when *A*_*i*_ = 2, where the biological effect sizes are zero and *b*^2^ is zero. Applying MPRAudit to the data, Figure 4a shows that *b*^2^ is close to zero and that the bias rapidly decreases as the number of sequences per clone increase, and that the calculated standard deviation agrees with the empirical truth.

**Figure 4:**
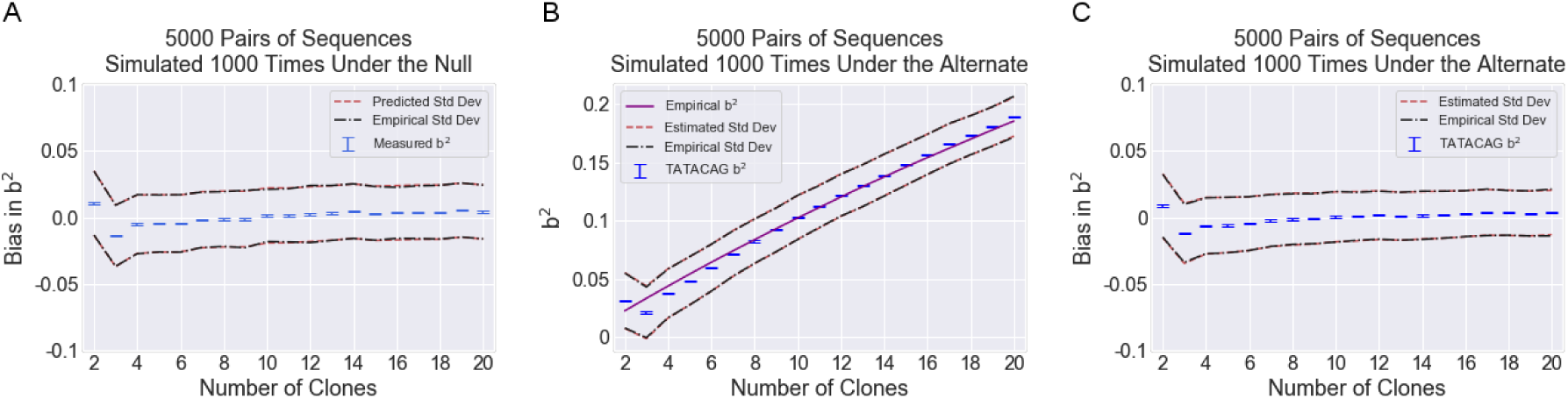
Results of explainability (b^2^) calculations for 5,000 pairs of sequences. The effect size in question is the random mutation of the TATACAG motif, which varies as a function of the number of T’s in the mutated sequence following equation (6). Error bars (very narrow on this scale) give the standard error of b^2^ for the 5,000 pairs of sequences. Dashed lines give estimated and empirical confidence intervals (standard deviation). (a) Effect size variability for pairs of sequences under the null, where b^2^=0. (b) Under the alternate, where the effect sizes vary in correspondence with the number of T’s in the mutated sequence. (c) Same as (b), except the y-axis gives the bias (estimate minus empirical).

Next, to introduce biological variability between WT-MT pairs, we apply MPRAudit to the same pairs of sequences under the model *A*_*i*_ = 2 − 0.5*f*_1*i*_ + 0.15*f*_2*i*_. If *f*_2*i*_ were constant, the change in gene expression between WT and MT would be uniform (Δ*A*_*i*_ = 0.5), and we would expect the explainability to be zero. With *f*_2*i*_ variable, the change in gene expression between WT and MT varies according to the number of Ts in the mutation, so we expect *b*^2^ to be nonzero. Figure 4b shows that this is what we observe in the data, and Figure 4c shows that the bias is uniformly small and decreasing with increasing clones per sequence. Overall, Figure 4 demonstrates that MPRAudit can determine the degree to which interesting biology has an impact on the effect size of mutations or variants in an MPRA.

In the above example we set the flanking sequence to be constant and randomized the mutation, but another interesting application of MPRAudit occurs when the flanking sequence varies and the mutation remains constant. Applying MPRAudit to the latter case will determine the importance of the flanking sequence to the effect of the mutation. This application might be important when considering the genetic context of binding sites and other sequence motifs.

## Real Data

### Overview

To demonstrate the utility and performance of MPRAudit in realistic settings, we looked at 12 real MPRA data sets from two different MPRA technologies. In each case we require at least 5 clones or barcodes per sequence. We also require a minimum number of counts per clone to filter out sequences with low statistical power and we follow the analysis guidelines of the published work when given.

The first several datasets are from our fast-UTR study of 3′ UTR transcripts in human cell lines. We analyze data from two cell lines (Jurkat and Beas2B)[4]. In the design of this study there are 41,288 distinct 3′ UTR sequences (including common variants and deliberate mutations) from 4,653 genes, which are each 160 nt long. Each sequence has twenty clones on average, which differ only by an 8 nt barcode. Not all sequences were observed in all sequencing runs. For this experiment we require each measured sequence to have at least five clones with more than five counts of DNA, and for at least one clone of each measured sequence to have at least one count of RNA. Any sequence that does not satisfy these conditions is excluded from the study. We then analyze sequences according to the statistic RNA/(RNA+DNA):

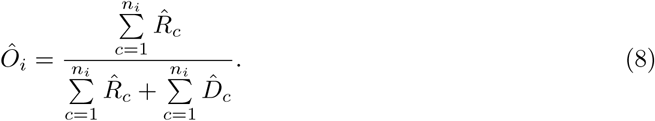

The study is designed to investigate the effects of several classes of active motifs and introduces mutations to determine the impact of those motifs [10, 17, 18, 35, 24] This study considers single sequences, pairs of sequences, and groups of sequences in the course of its investigation.

The second dataset is from a publicly available MPRA[23] where the authors compare the functional activities of 2,236 candidate liver enhancers in episomal (EPI) and chromosomally (CHR) integrated contexts. Three replicates of each experiment were made available to the public online, so we have compared the datasets individually and to each other according to the statistic used in their study:

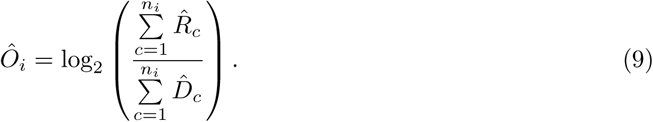

For this dataset we did not place a filter on the minimum number of counts per barcode, as the smallest number is already seven. We combine the replicates at the clonal level by first normalizing each replicate by dividing by the total number of counts of RNA or DNA, then combining replicates by concatenating the normalized data files. Thus, the combined dataset effectively behaves as if it has a greater number of measured clones.

### Single Perturbations: Contribution of individual sequences to variation in outcomes

#### Experiment-wide measures of noise

For each data set we estiamte the total *b*^2^ and provide all results in Table 1. For the fast-UTR data, the results show that a large fraction of the variance in these datasets (≈50%) is explained by sequence variation, which means accurate predictors of gene expression should be possible to generate for these datasets. However the noise might be too high to establish accurate estimates of activity at the single-sequence level, and deeper sequencing could improve the signal. For the Inoue data sets, the *b*^2^ is approximately 90%, implying a very low contribution from technical noise and only a modest benefit could be obtained form deeper sequencing.

**Table 1:**
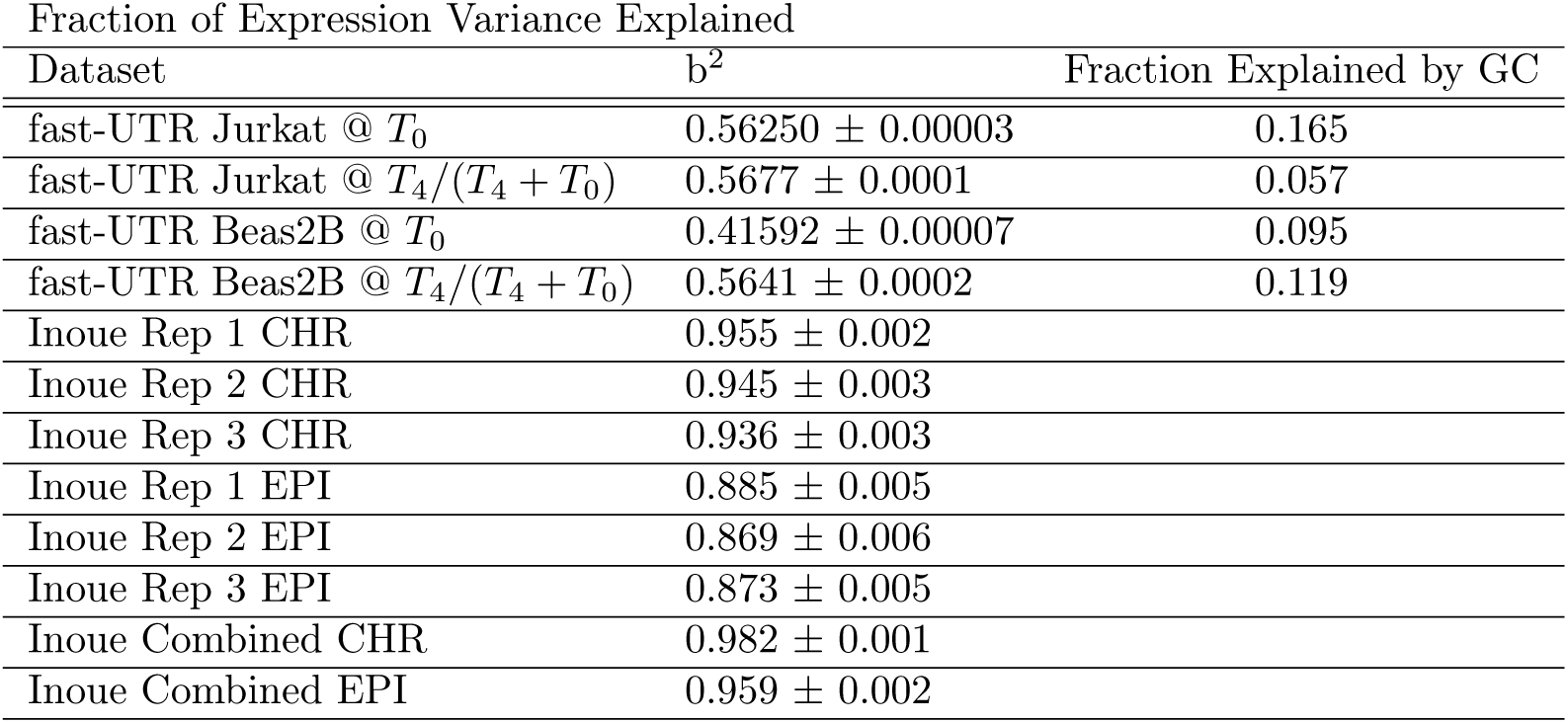
Fraction of Variance Explained by Sequence for Several Datasets, and Fraction of Variance Explained by GC for Fast-UTR data.

#### GC-content biases aggregated outcomes

We also estimated the fraction of variation caused by GC content alone, by residualizing at the sequence level and determining the change in variance. To do this we fit a 5th-order polynomial to the RNA/(RNA+DNA) ratio as a function of GC-content and subtract it from the gene expression (see supplement for further details). In general, we find that more GC leads to a decrease in gene expression, which runs counter to the trend of the AU-rich elements that this dataset is enriched for, indicating that GC-content is a separate effect from AU-mediated decay.

#### Performance of computational prediction methods

Several methods have been proposed to predict the outcome of MPRA experiments directly from sequence[25, 26, 27]. The performance (squared correlation between predicted and measured values) is bounded above by *b*^2^. We trained a k-mer lasso model[26] in the fast-UTR data set, which takes sequence as its input, and outputs a prediction of the RNA/DNA ratio (see Methods). The squared-correlation between the out of sample prediction and measured outcome was at most 0.26. Comparing these percentages to Table 1, we expect that this approach can be substantially improved with better methods and/or less noisy data. Similarly, in the Inoue data sets, we find that the *b*^2^ of the datasets are much higher than the squared correlation of their prediction methods (Pearson *R*^2^ of 0.34 on sequence data and 0.36 when combining chromatin annotations with sequence information), which indicates that most of the sources of sequence-related variation remain unexplained by the current set of predictors.

### Groups of Perturbations: How much variation is unaccounted for by existing biological categories?

#### Assessing the contribution of CDE stem length to 3’UTR function

From the fast-UTR data, we consider an example involving the stem lengths of constitutive decay elements (CDEs) in the 3′ UTR. CDEs are conserved stem loop motifs that the proteins Roquin and Roquin2 bind to in order to promote mRNA decay.[36] We classify CDEs of the form UUCYRYGAA as having a stem length of 2-5 nt if the surrounding 2-5 nt form reverse-complementary pairs (e.g. CCUUCYRYGAAGG has a stem length of 2). We observe this motif to be destabilizing, and we also observe that increasing the stem length increases the decay rate: linear regression on the decay of gene expression with stem length produces a slope that differs from zero with *p* = 7.5 × 10^*−*7^, *r*^2^ = 0.10, and *slope* = −0.063.

Using MPRAudit, we can examine the full set of sequences with CDEs and determine that *b*^2^ = 0.8598 ± 0.0002, so 86% of the total variance is determined by differences in sequence. Accounting for categories of CDE stem lengths, MPRAudit determines that the within-group explainability is 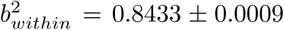. The fraction of within-group variation explained by sequence variation has barely changed as a result of grouping by CDE stem length, because the number of sequences affected by these groupings are small in number. We conclude that stem length has a statistically significant effect, but accounts for only a small fraction of the variation in CDE containing sequences.

We can also look within each category of CDE stem length and examine the *b*^2^ for each length of stem (2-5 nt). Table 2 shows that the variation due to sequence within groups 2 and 3 are roughly the same as the whole, and since they make up the majority of sequences, they dominate the total within-group explainability. On the other hand, the b^2^ within groups 4 and 5 drop progressively, with the smallest amount of sequence-based variance in group 5. Table 2 also specifies which groups of sequences warrant further study. Within groups of fixed CDE stem length, the greatest variation occurs when the stem length is 3, and the least occurs when the stem length is 5. This suggests that more novel biology is likely to be uncovered within the former group of sequences than the latter, even though the latter have the more active elements. The fact that *b*^2^ happens to be much higher within these particular groups than the dataset average given in Table 1 suggests that these are particularly active regions and that there are further rules for CDE activity that are waiting to be uncovered.

**Table 2:** Explainability of sequences with CDEs, classified by stem length

| Stem Length (nt) | $b^2$ | $N_{Sequences}$ |
| --- | --- | --- |
| 2 | $0.8564 \pm 0.0006$ | 139 |
| 3 | $0.8651 \pm 0.0007$ | 71 |
| 4 | $0.7983 \pm 0.0019$ | 29 |
| 5 | $0.6561 \pm 0.0309$ | 7 |

### Pairs of Perturbations: What is the contribution of designed mutations to variation in measured outcomes?

In the fast-UTR data we define the effect size of a mutation to be the signed difference in gene expression between a sequence and its mutant. To estimate the variance contributed from the mutations we run MPRAudit over the signed differences of paired sequences. If the mutations have no effect then the estimate of *b*^2^ will be 0. Of the 41,288 distinctly designed sequences in the dataset, 17,066 sequences are deliberately designed mutations of a known motif, which are present in 9,754 controls (many of the controls have multiple mutations).

The results of this analysis are presented in Table 3. Interestingly, the mutations in the fast-UTR contribute primarily to the *T*_0_ timepoint in the Jurkat cell line and primarily to the ratio of *T*_4_*/*(*T*_4_ + *T*_0_) in the Beas2B cell line. We conclude that bottom-up approaches to identify specific mutational effects should focus on these subsets of the data.

**Table 3:**

| Data Set | Fraction of Variance Explained by Mutations |
| --- | --- |
| fast-UTR Jurkat @ $T_0$ | $0.1698 \pm 0.0002$ |
| fast-UTR Jurkat @ $T_4/(T_4 + T_0)$ | $0.093 \pm 0.001$ |
| fast-UTR Beas2B @ $T_0$ | $0.0103 \pm 0.0008$ |
| fast-UTR Beas2B @ $T_4/(T_4 + T_0)$ | $0.156 \pm 0.0004$ |
| Inoue UTR Rep 1 | $0.885 \pm 0.005$ |
| Inoue UTR Rep 2 | $0.869 \pm 0.006$ |
| Inoue UTR Rep 3 | $0.873 \pm 0.005$ |
| Inoue UTR Combined | $0.918 \pm 0.004$ |

We perform a similar analysis on the Inoue et al data on the difference in gene expression between episomal and chromosomally integrated experiments, and find that *b*^2^ remains high (≈0.9). The high value of *b*^2^ indicates that the differences in sequence effects between the episomal and chromosally integrated experiments are driven by currently unknown features that might be worth investigating in a future analysis.

Note that the results in Table 3 and Table 1 are not directly comparable. The results in Table 3 can be very large even if the results in Table 1 are close to 0 or vice versa. The effects of GC content matters less in this case due to cancellation of per-nucleotide GC, so we did not repeat our GC-content analysis.

## Discussion

In this work we demonstrate a method to determine the fraction of variance in a molecular assay that is due to sequence variation as opposed to technical artifacts or statistical noise. The purpose of this method is to help researchers who use MPRAs to know where to target their analysis efforts in different types of datasets as well as characterize the results of their existing analyses. Using realistic simulations of two types of MPRA data, we showed that MPRAudit is unbiased and calibrated provided a modest minimum clonal and sequencing depth is achieved. Application to real data demonstrates how MPRAudit can determine additional biological and experimental information about the sources of variation within a dataset.

In concurrent work [37], MPRAudit helped us to discover novel rules for sequence motifs. Table 1 in Ref. [37] shows that mutations of established classes of AU-rich elements (AREs) have a sizeable amount of explainability, ranging from *b*^2^ = 0.1 to 0.5 (depending on the category and time point). This indicates that the effect sizes of those mutations (and possibly the effect of flanking sequences on those mutations) varies significantly within the previously established categories. Suspecting that this classification system might lack important information, we were able to develop better rules for predicting the effects of AREs on gene expression in humans, improving on the existing classification by up to 50%.

Even if no single mutation is statistically significant at the level of the study, MPRAudit can be used to identify candidate groups or sets of sequences with high explainability.[38] Applications may be found in relation to pathway analysis or genes or annotations of interest.

MPRAudit does not rely on parametric model assumptions and is broadly flexible with respect to the quantity being measured. We rely on jackknife resampling rather than some sort of weighted linear model because the paired counts of RNA and DNA make it difficult to calculate the proper weights for each clone, whereas this information is automatically incorporated in the jackknife.

MPRAudit relies on only a single replicate experiment, which mitigates confounding batch effects, saves time and money, and is better at handling sparse datasets than measuring correlations between replicates. In the results section we note that there is a relationship between the correlation between replicates and the explainability: the squared correlation between replicates should approximately equal the product of the explainabilities; a short proof is given in the supplement. If the correlation between two replicates is low, the correlation between them is not capable of determining whether one replicate is less reproducible than the other.

It should be kept in mind that the correlation between signal and noise may be nonzero when the counts of RNA or DNA are very low, leading to nonsensical explainability estimates and the breakdown of our additive model in equation 2. It should also be noted that since the technical and total variances are estimated separately, when the explainability is truly zero the estimate of the explainability intuitively should be negative for roughly half of all measurements.

Resampling is most effective when the number of samples is large. This can be a problem for both the number of clones per sequence for *V ar*(*ϵ*), and the number of sequences in the dataset for *V ar*(*Ô*). To address the former issue, we suggest placing a filter on the minimum number of clones. In our analysis we required at least 5 clones per sequence, but a higher cutoff would give greater accuracy. With regards to the number of sequences in the dataset, we acknowledge that MPRAudit performs best on a massive dataset such as MPRAs.

Sparse datasets can also cause problems with subsampling or resampling, since each of the jackknifed subsamples must be well-defined. If *O*_*i*_ is given by 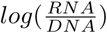, each jackknifed subsample must have a nonzero number of RNA counts and DNA counts. For sparse datasets a statistic such as 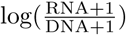 might be worth considering. This will not be an issue for 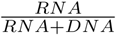, which is well-defined as long as the total counts are nonzero, but it might become a problem in a ratio of ratios such as. 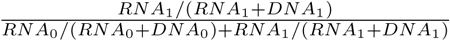.

Missing data is also an important quality control issue. While having very few counts per clone is not in theory a limitation of MPRAudit as long as the number of clones is high, we find that requiring at least one count of RNA and DNA per sequence gives us far greater repeatability across replicate datasets in practice due to dropout[39]; the set of missing sequences has a low correlation between replicates. These cases produce erroneously low predictions of technical variance, since resampling will repeatedly produce the same 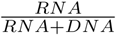 ratio, predicting zero technical variance while creating the largest possible amount of variance in practice.

The convergence and bias of various resampling techniques will differ for different methods. For instance, in our analysis of simulated data, using *log*(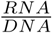 led to greater bias than 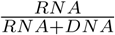. These considerations should be weighed by the user and investigated for their particular use case. Since MPRAudit estimates the variance of the noise of each measurement, we anticipate that these techniques will be directly applicable to many related methods that can be applied to MPRAs. For example, sampling-based variances can be used to provide the weights to regressions, ANOVAs, Bayesian analyses[14], and other methods that accept precision weights in heteroskedastic models, improving the power of bottom-up MPRA analyses.

The code that we have made freely and easily accessible gives some implementations of MPRAudit, and can be easily extended by altering just a few lines. Users can make extensions of MPRAudit to consider arbitrary choices of statistics, such as complex ratios of ratios or regressions of time series data. Users can also swap the delete-D Jackknife with other resampling techniques.

## Methods

### Introduction

MPRAudit allows us to estimate the fraction of the total outcome variance due to the variation from clone to clone. At the sequence level, we measure a collection of N sequences *i* ∈ {1…*N*} with some outcome *O*_*i*_:

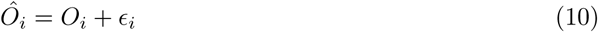

where *ϵ*_*i*_ is the error in the measurement, and we make no assumptions about the distribution of *E*_*i*_. We measure *Ô*_*i*_ directly, but we cannot know the values of *O*_*i*_ or *ϵ*_*i*_. However, we can estimate *V ar*(*ϵ*_*i*_|*i*) using the method we establish below. From there we build up methods to estimate the explainability b^2^ and the uncertainty of the explainability 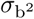.

### Calculation of *V ar*(*ϵ*_*i*_|*i*)

The workhorse of MPRAudit is the calculation of *V ar*(*ϵ*_*i*_|*i*).

First note that for clones of a single sequence,

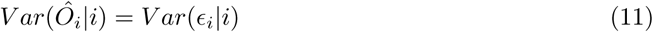

This variance can be estimated by resampling over the barcoded clones of a sequence, which we do with a delete-d-clone jackknife:

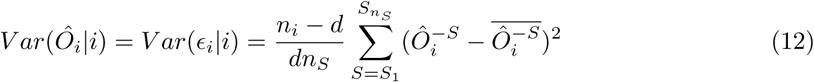

where 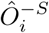 leaves out the S^th^ set of d clones, and there are 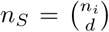 subsets of d clones in *n*_*i*_ total clones. So if we use

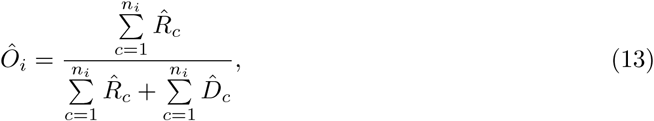

then

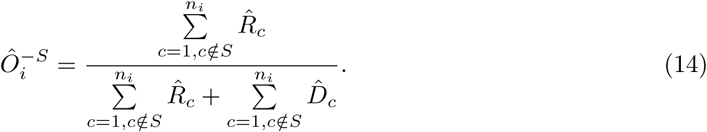

Or if

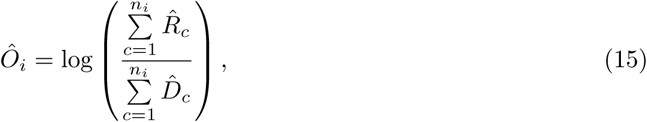

then

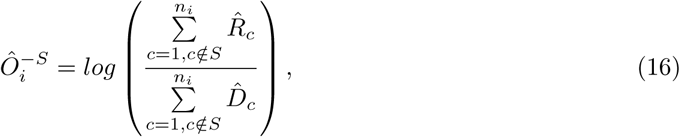

and so on. According to the literature[34], the optimal value of d is in the range 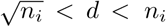, which can make the total number *n*_*S*_ of subsets very large when *n*_*i*_ is of modest size and *d* differs from *n*_*i*_. Since it is computationally intractable to sum over all possible subsets, as a first implementation we use 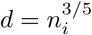 and sample over random subsets.

There are several types of bias-corrected bootstrap and jackknife techniques (including delete-D jackknifes) in the literature[33, 40, 34, 41, 42], and it is beyond the scope of this paper to explore the entire space of possible resampling techniques and parameters. On null data, we observe that the delete-one jackknife tends to underestimate our values of *b*^2^, whereas the standard bootstrap tends to overestimate it (and the bias of the bootstrap is larger than the jackknife). We use the sampled delete-D jackknife because it appears to have less bias for sequences with smaller numbers of clones.

### Fraction of Variance Explained by Sequence Variation

We would like to be able to estimate the fraction of variance explained by differences in sequence, which we define as as:

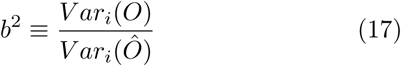

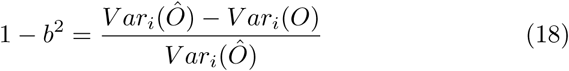

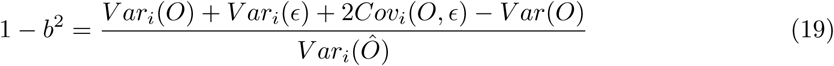

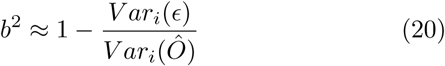

where we neglect the covariance term as being small. For *V ar*_*i*_(*ϵ*), we use

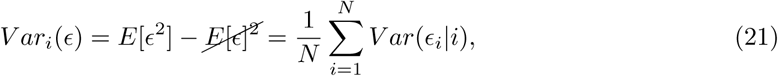

where we assume that the error is unbiased, and *V ar*(*ϵ*_*i*_) is the quantity estimated by MPRAudit in equation 12.

### Fraction of Variance Within Groups Explained by Sequence Variation

We are interested in determining the fraction of variance within K groups of sequences that is due to sequence variation. Let n_*j*_ be the number of sequences within group j (*j* ∈ {1…*K*}), let *Ôij* be the value of sequence i in group j, let 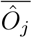 be the sequence average within group j, let 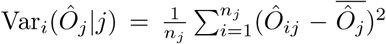 be the variance across sequences within group j, and let 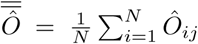 be the grand mean over all sequences in all groups. Then without loss of generality, one can write:

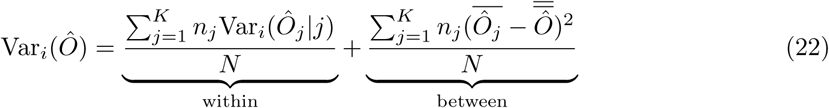

The left hand side of equation (22) is the variance over all sequences. On the right hand side, the first term is only a function of the variances within groups, and the second term is only a function of the means within groups and the grand mean. Therefore the first term on the right is the contribution of the within-group variation to the total variance, and the second term on the right is the contribution of the between-group variation.

Since the within-group variance is just a sum of variances and the measurements are assumed to be independent, the variance due to clonal variation can be calculated following the method of obtaining *ϵ* in section 4.1.2: the clonal variance *ϵ*_*j*_ of each group j can be computed for each *V ar*_*i*_(*O*_*j*_ |*j*), then added according to (22):

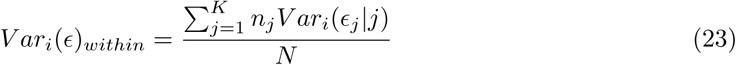

and

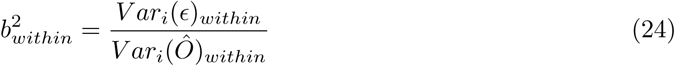

### Fraction of Variance Explained by Pairs of Perturbations

Considering pairs of sequences for an effect size analysis follows a similar framework as above. Given a collection of pairs {i,j} of sequences with some outcome *O*_*ij*_, we define the measured effect size of a pair as:

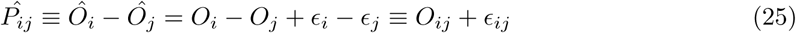

Following the same logic as above, we obtain

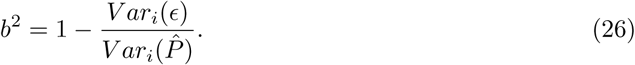

where

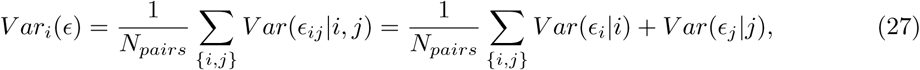

if we assume the covariance is small, where *V ar*(*ϵ*_*i*_|*i*) is directly calculated by MPRAudit as above.

### Estimate of Uncertainty

The uncertainty of these estimates of technical variance can be computed by jackknifing again (here we use the standard leave-one-out jackknife). For the uncertainty of the estimate of the error, we use the jackknife estimate of the variance of the mean of the jackknife estimates:

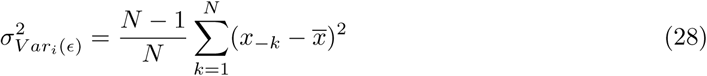

where 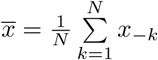 and

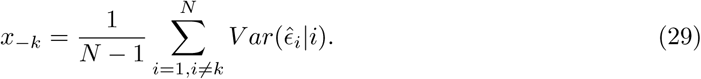

For the total observed variance we use the jackknife estimate of the variance over outcomes:

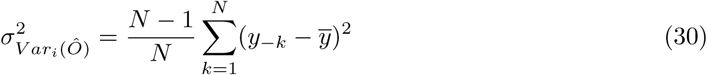

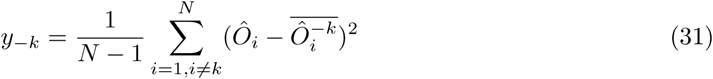

where 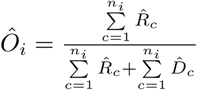 as above, 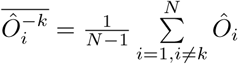 and 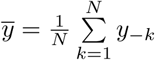.

This gives the single-perturbation case, and can be trivially extended to pairs and within groups of perturbations as above.

### Estimate of Fractional Uncertainty

In the previous section we estimate the uncertainty of *V ar*_*i*_(*ϵ*) and *V ar*_*i*_(*Ô*), but we mostly care about the uncertainty of the ratio 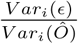. Given a ratio of the form 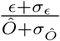, the first order result in *σ*_*ϵ*_ and *σ* _*Ô*_ is 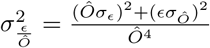 (again neglecting the covariance term), and 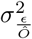 is the uncertainty of b^2^.

## Availability

Method and code are available for download at https://github.com/david-a-siegel/MPRAudit

## Acknowledgments

DAS, AB and NZ were funded by NIH grants K25HL121295, U01HG009080, R01HG006399, R01CA227237, R03DE025665, R01CA227466, R01ES029929, R01GM110251, R01HL124285, and DoD grant W81XWH-16-2-0018

DJE and OLT were funded by NIH grants R35 HL145235, R01 GM110251, and R01 HL124285. DAS would like to thank Andrew Dahl, Joel Mefford, and Nadav Ahituv for useful discussions.

