## Supplement for "MPRAudit Quantifies the Fraction of Variance Described by Unknown Features in Massively Parallel Reporter Assays"

#### Details of the Analysis of Real Data

As stated in the main text, the fast-UTR dataset had two distinct cell lines. The data from each cell line were obtained in two sequencing runs, which we then combined to form higher statistic datasets.

To combine the Jurkat data, we first normalized the DNA counts (divided by an overall constant) to set the total RNA:DNA count ratio to be equal in both sequencing runs. We prefer this to other methods because 1. Simple addition would create a bimodal distribution of RNA/DNA ratios; and 2. Normalizing both the RNA and DNA counts (by dividing by the total number of counts) treats each sequencing run as equal, whereas one sequencing run had more counts and clones than the other.

For the Beas2B data we did not have counts for both RNA and DNA from both sequencing runs so we simply added the counts directly.

#### GC Content

To account for GC-content in the fast-UTR data, we fit a 5th-order polynomial to the measured outcomes as a function of GC-content and subtract it from the measured outcomes. This decision was made by assessing the performance of Nth order polynomial fits ($N \in \{1, ..., 7\}$) through leave-one-out cross validation. After iterating over the dataset, we compare the predicted outcomes of the left out sequences to their actual outcomes by computing the Pearson correlation. The results for the Jurkat data are given in Table 1. Since the correlation was maximized at N=5 for the $\frac{T_4}{T_4+T_0}$ data and only increased slightly after N=5 for the $T_0$ data, we decided to use N=5. The 5th order polynomial fits are shown in Figure 1.

| N | LOOCV Corr. at $T_0$ | LOOCV Corr. at $T_4/(T_4+T_0)$ |
|---|---|---|
| 1 | 0.3660 | 0.1137 |
| 2 | 0.3820 | 0.1417 |
| 3 | 0.3825 | 0.1705 |
| 4 | 0.3839 | 0.1708 |
| 5 | 0.3840 | 0.1736 |
| 6 | 0.3843 | 0.1736 |
| 7 | 0.3843 | 0.1734 |

Table 1: Correlation between predicted and actual gene expression for Nth order polynomials for data measured at $T_0$ and $\frac{T_4}{T_4+T_0}$.

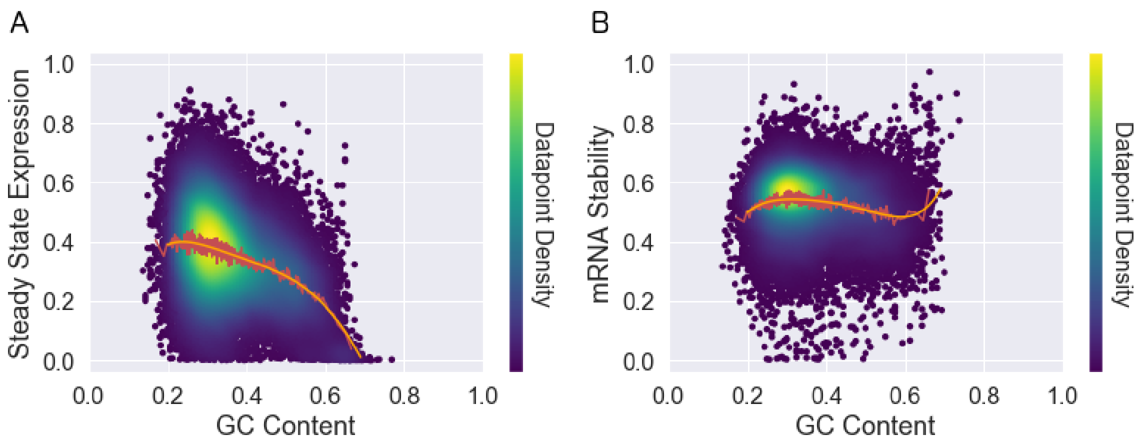


Figure 1: Gene expression and decay rates vary as a function of GC content at each time point. We focus on $T_0$ (steady state expression) and $\frac{T_4}{T_4+T_0}$ (mRNA stability). Each dot is one sequence, with a color scale given by the density of surrounding data points calculated with a Gaussian kernel and the highest-density points plotted last. The red line is an 80-sequence moving average; the orange line is the 5th-order polynomial fit that we use to residualize the data.

### Further Details of Our Simulation

Once simulated populations of RNA and DNA molecules are generated, RNA and DNA counts for each clone and sequence are drawn from a negative binomial distribution in order to simulate overdispersion. The means are proportional to the true number of simulated molecules and to the total number of counts:

$$\hat{R}_{ic} \sim NBinom(\mu_R, \sigma_R^2)$$
$$\hat{D}_{ic} \sim NBinom(\mu_D, \sigma_D^2)$$
where
$$\mu_R = K_R \times R_{ic}$$
$$\mu_D = K_D \times D_{ic}$$
$$\sigma_R^2 = P \times \mu_R$$
$$\sigma_D^2 = P \times \mu_D$$

where $K_R$ and $K_D$ are normalization constants (the ratios of the expected counts of RNA and DNA to the number of RNA and DNA molecules, respectively), and P is a constant greater than one (and typically much larger than one) that allows us to adjust the overdispersion of the negative binomial.

To simulate data we used the following parameters: $\lambda = 20$, $s = 0.1$, $P = 30$, $K_R = 1.5$,$K_D = 1$. This leaves only the choice of $A_i$ to be varied within an experiment, which is convenient since $A_i$ is an intrinsic quantity (the expected RNA/DNA ratio doesn't change with sequencing, the number of DNA, the amount of noise, and so on). For simulations with no biological variation, $A_i$ is set to be constant and we used $A_i = 2$.

Since $K_R$ and $K_D$ are proportional to the total counts of RNA and DNA, increasing their values decreases the fraction of variance explained by $\sigma_R^2$ and $\sigma_D^2$. We set $K_R$ to be slightly larger than $K_D$ simply to avoid cancellation by symmetry. These parameters (including $A_i = 2$) correspond to an average of approximately 50 and 20 counts of RNA and DNA per clone, respectively.

$P$ is the parameter that sets the amount of overdispersion, which is common in sequencing experiments.[1] If P is set to one, then the mean becomes equal to the variance as in a Poisson distribution.

### $b^2$ is the Upper Limit to the Squared Correlation Between Prediction and Data

Say you have a real dataset:

$$\hat{O}_i = O_i + \epsilon_i$$
$$i \in \{1...N\}$$

and a perfect prediction method:

$$O_{pred} = O$$

Then the correlation between the data and prediction is:

$$corr_i(\hat{O}, O_{pred}) = \frac{cov_i(\hat{O}, O_{pred})}{\sqrt{var_i(\hat{O})var_i(O_{pred})}}$$
$$cov_i(\hat{O}, O_{pred}) = cov_i(O + \epsilon, O) = var_i(O) + \cancel{cov_i(\epsilon, O)}$$
$$corr_i^2(\hat{O}, O_{pred}) = \frac{var_i^{\cancel{2}}(O)}{var_i(\hat{O})\cancel{var_i(O_{pred})}} = b^2$$

If the prediction method is less than perfect then the squared correlation will be lower. So the explainability $b^2$ provides an upper limit to the performance of a prediction method.

### Correlation Between Replicates

There is a relationship between the explainability and the correlation between replicate datasets:

$$\hat{O}_i^a = O_i + \epsilon_i^a$$
$$\hat{O}_i^b = O_i + \epsilon_i^b$$
$$i \in \{1...N\}$$

$$corr_i(\hat{O}^a, \hat{O}^b) = \frac{cov_i(\hat{O}^a, \hat{O}^b)}{\sqrt{var_i(\hat{O}^a)var_i(\hat{O}^b)}}$$
$$= \frac{var_i(O) + cov_i(O, \epsilon^a) + cov_i(O, \epsilon^b) + cov_i(\epsilon^a, \epsilon^b)}{\sqrt{var_i(\hat{O}^a)var_i(\hat{O}^b)}}$$
$$\approx \frac{var_i(O)}{\sqrt{var_i(\hat{O}^a)var_i(\hat{O}^b)}}$$
$$= \sqrt{b_a^2 b_b^2}$$

This is close to what we measure in the Inoue et al [2] datasets.

### Additional Method Simulations

To further demonstrate the conditions under which MPRAudit is unbiased and that our simulation works as we expect it to, we have simulated data under a wide range of parameters, which we show in figure 2. Figure 2a shows that MPRAudit performs accurately even when the value of "A", the true RNA/DNA ratio, is extreme in comparison with the average. This shows that extreme values of RNA/(RNA+DNA) do not lead to biased estimates of $b^2$. When the RNA/DNA ratio is very low there is some deviation between the estimated and empirical standard deviation of $b^2$, due to the very low number of RNA counts in that regime.

Figure 2b shows the effect of the number of counts on a dataset with no signal. There is some bias when the number of counts is very low (when there are less than roughly ten counts per clone), but the bias quickly drops to less than a few percent when the number of counts is higher than

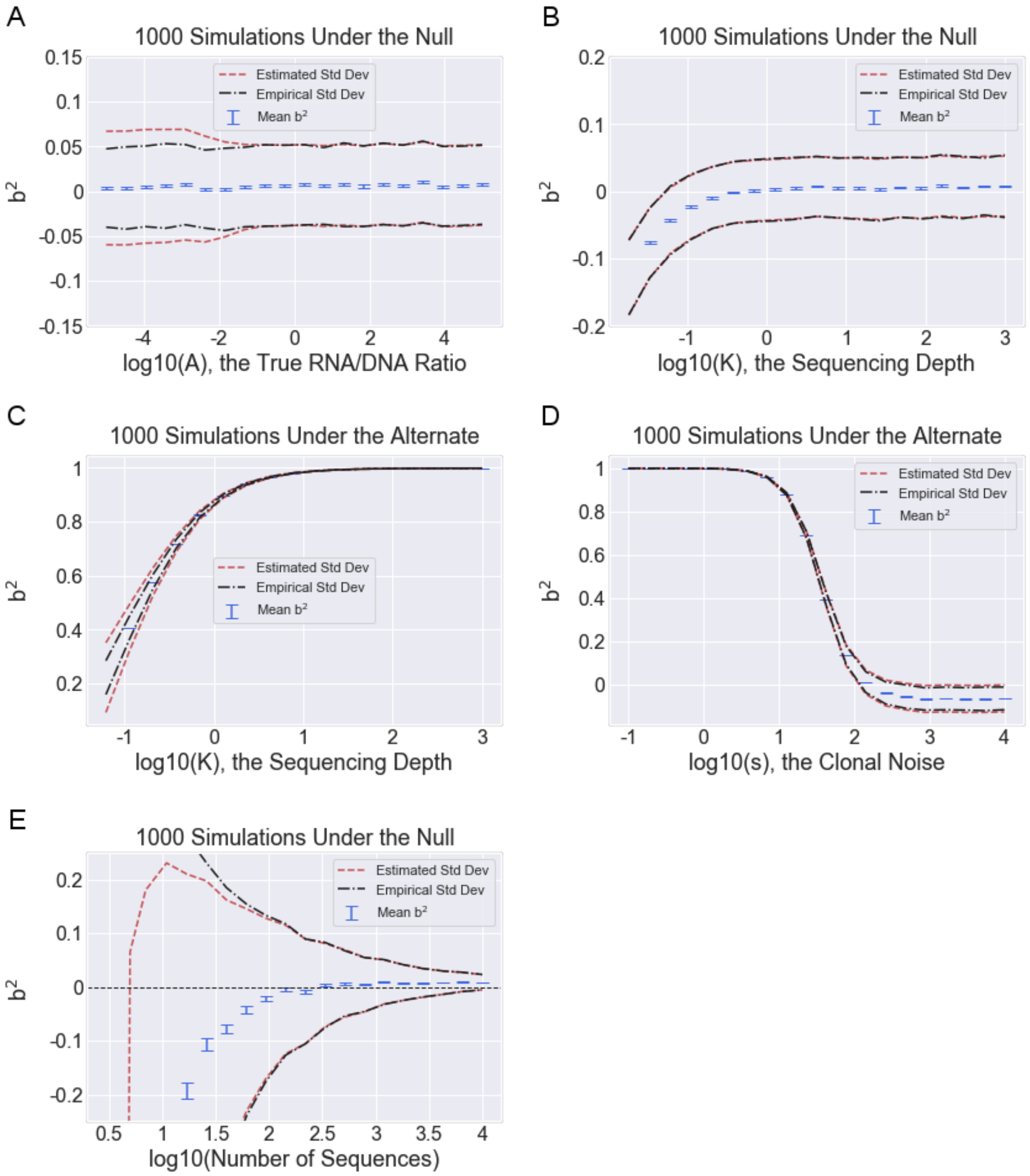


Figure 2: (a) Performance of MPRAudit for different values of $A$ under the null. Parameters for these simulations: 1000 sequences, 100 clones, 20 DNA/clone, $s = 0.1$, $P = 30$, $K = 10$. (b) Performance of MPRAudit for different values of $K$ under the null. Parameters for these simulations: 1000 sequences, 100 clones, 20 DNA/clone, $A = 2$, $s = 0.1$, $P = 30$. (c) Behavior of MPRAudit under the alternate for different values of $K$, the sequencing depth. Parameters for these simulations: 1000 sequences, 100 clones, 20 DNA/clone, $A = |N(0, 0.1)|$, $s = 0.1$, $P = 30$. (d) Behavior of MPRAudit under the alternate for different values of $s$, the clonal noise. Parameters for these simulations: 1000 sequences, 100 clones, 20 DNA/clone, $A = |N(0, 0.1)|$, $P = 30$, $K = 10$ (e) Performance of MPRAudit for different numbers of sequences. Parameters for this simulation: 1000 sequences, $100 \times \textit{ratio}$ clones, 20 DNA/clone, $A = |N(0, .01)|$, $s = 0.1$, $P = 30$, $K = 10$.

roughly ten per clone. For this set of parameters, when K=1 ($log_{10}(K) = 0$) we have approximately 50 counts of RNA and 20 counts of DNA.

Figure 2c shows the effect of increasing the number of counts on a dataset with sequence-based variation. The sequence-based variation is set to remain constant throughout (we draw the value of A, the sequence RNA/DNA ratio, from a random normal with mean zero, variance 1, and we use the absolute value). When the number of counts are low, the explainability is low due to sequencing-based noise, and the empirical standard deviation differs slightly from the estimated standard deviation. As the number of counts increases the estimated and empirical standard deviations match, and the explainability increases.

Figure 2d shows the effect of increasing the clonal noise on the dataset. When the clonal noise is low, the explainability is very high. As the noise increases, $b^2$ decreases until the noise dominates the signal.

Figure 2e shows the importance of having a large number of sequences in the dataset. Under these conditions, MPRAudit works best when the number of sequences is large.
